# Biodegradation of components from an oxidized polyethylene by a *Rhodococcus* strain isolated from the gut of Atlantic Salmon

**DOI:** 10.64898/2026.03.19.712908

**Authors:** Ronja Marlonsdotter Sandholm, Dave Rojas Calderon, Marcus Torres Hansen, Ravindra R. Chowreddy, Gustav Vaaje-Kolstad, Sabina Leanti La Rosa

## Abstract

Polyethylene (PE) is the most produced synthetic polymer and as a result, a major source of microplastic waste accumulating globally. Exposure to photo- and thermo-oxidative conditions in the environment can cause PE to degrade into carbonyl-containing compounds, hydrocarbons, and low molecular weight PE (LMWPE). In both marine and freshwater ecosystems, fish, including Atlantic salmon, can ingest PE and its derivatives, creating opportunities for interactions with their gut microbes. Here, we investigated the ability of a bacterial isolate from the gut of salmon, *Rhodococcus* sp002259485 strain ASF-10, to grow on an LMWPE model substrate for partially depolymerized and oxidized PE. Comparative genomic analyses showed that ASF-10 has a smaller genome than other *Rhodococcus* species yet retaining conserved functions including those related to utilization of medium- and long-chain hydrocarbons. In-depth characterization of the substrate following growth with ASF-10 confirmed depletion of alkanes and 2-ketones deriving from LMWPE, while the polymeric component remained unchanged. Proteomic analysis identified multiple enzymes that were likely to be involved in the degradation of LMWPE-derivatives, including an alkane 1-monooxygenase, cytochrome P450 hydroxylases and Baeyer-Villiger monooxygenases, as well as proteins for production of biofilm and a surfactant that may enhance accessibility to the substrate. Collectively, our findings advance the understanding of the ecology and enzymatic mechanisms underlying utilization of medium- to long-chain alkanes and oxidized variants thereof, that resemble molecules that can occur from abiotic PE degradation, by a fish gut-associated microbe. This metabolic capacity could be harnessed to develop sustainable strategies for bioremediation of oxidized, LMWPE-derivatives.

**Importance:** The widespread presence of plastics in marine and freshwater environments has raised concerns due to their toxicity when ingested by fish. Microbial mechanisms driving breakdown of microplastic components, such as LMWPE and derivatives, in gut systems remain poorly understood. This study reveals how a bacterium isolated from the gut of salmon, *Rhodococcus* sp002259485 strain ASF-10, metabolizes alkanes and oxidized variants thereof, that can result from abiotic PE decomposition. We identified key enzymes that are potentially involved in this process as well as in the production of biofilm and surfactants that may facilitate access to the substrate. Besides extending the knowledge of the enzymatic basis for degradation of PE-derivatives in gut-associated microbes from aquatic organisms, our results provide a framework that couples advanced compositional characterization of the substrate with omics techniques, offering valuable insight to support future studies aimed at unequivocally identifying microbes and their enzymes implicated in transformation of PE-derivatives.

## Introduction

Persistent plastic and microplastics (MPs) pollution represents a novel niche within the aquatic environment, serving as an attractive substrate for bacterial growth or a favorable surface for microbial colonization and biofilm development in the otherwise open-water systems (1). In recent years, a multitude of studies have explored the marine “plastisphere”, detailing the diversity, composition and antibiotic profile of the microbial community colonizing plastic surfaces (2, 3). Aquatic animals, including wild and farmed Atlantic salmon (*Salmo salar*), are exposed to MPs through multiple pathways, including uptake via gills, ingestion of contaminated prey, or consumption of polluted feed (4). Additionally, the aquaculture industry relies heavily on plastic materials for both infrastructure and feed packaging. These include polyethylene (PE), polyethylene terephthalate (PET) and polypropylene (PP) that are used to make cages, ropes, cords, and fishing nets, as well as polystyrene (PS) in foam-filled collars for fish cages, polymer-coated nets, plastic bins for harvesting, feed sacks, and foam buoys. In salmon farming, cages and tanks, as well as feeding systems, are primarily built of high-density PE (HDPE; molecular weight of 300,000-500,000 g/mol), while low-density PE (LDPE; molecular weight of 50,000-300,000 g/mol) is used in protective coatings (5–7). UV light, high temperatures or oxidizing agents can lead to polymer oxidation and depolymerization, resulting in formation of low molecular weight PE (LMWPE; molecular weight of 1,000 – 50,000 g/mol; chain length of 70-1400 carbon atoms) as well as oxidized and non-oxidized fragments, such as ketones and alkanes with a chain length of 5-36 carbons (8, 9). While it is reasonable to assume that PE and PE-derivatives may leak and be ingested by the fish, no studies to date have examined how gut microbes from aquatic animals could actively interact with MP-derived components. This highlights a critical knowledge gap and underscores the need for further research to determine the fate of PE and its derivatives within the salmon digestive system, including characterization of the metabolic potential and expressed functions of microorganisms interacting with these compounds.

Our group has recently combined targeted isolation strategies with state-of-the-art whole genome sequencing to generate the Salmon Microbial Genome Atlas (SMGA), a culture collection of bacterial isolates from Atlantic salmon with fully sequenced genomes (10). Additional isolation efforts have genotypically characterized a strain taxonomically assigned as *Rhodococcus* sp002259485 strain ASF-10 (hereafter referred to as *Rhodococcus* sp. ASF-10) (11). Access to these well-characterized strains supports research into how salmon gut bacteria process and utilize key nutrients such as proteins, lipids, and carbohydrate from feed, or possibly interact with “uncommon” polymers in the gut, such as MPs and hydrocarbons.

In this study, using a combination of bioinformatic approaches, proteomics and analytical techniques, we unravel the molecular mechanism deployed by *Rhodococcus* sp. ASF-10 for the degradation of an oxidized, low molecular weight PE (LMWPE) that resembles an abiotically oxidized form of polymeric PE. We provide genomic evidence demonstrating conserved metabolic functions and the presence of genes encoding enzymes linked to alkane degradation in *Rhodococcus* sp. ASF-10 as well as in a multitude of other *Rhodococcus* isolates from different environments, highlighting the genus’s ability to mineralize these compounds. Proteomic analysis and in-depth substrate characterization demonstrated that *Rhodococcus* sp. ASF-10 actively metabolizes derivatives from oxidized LMWPE, while did not utilize the polymeric fraction. Finally, network analysis of the proteomics data showed strong correlations between the consumption of the LMWPE derivatives, the pathways for alkane degradation, uptake and β-oxidation of fatty acids, as well as the production of biofilm and biosurfactants.

## Materials and methods

### Bacterial strain and growth conditions

*Rhodococcus* sp. ASF-10 was isolated from the gut of Atlantic salmon (*Salmo salar*) fry raised in freshwater conditions (11). The strain was routinely cultivated in Tryptone Soy Broth (TSB, Thermo Scientific, CM01219) at 22°C without agitation. Growth experiments were carried out in minimal salts medium (MM) supplemented with 30 mg/mL of either oxidized low molecular weight PE (LMWPE, M_w_ of ∼4000 g/mol, Sigma-Aldrich, 427772), sodium succinate (Sigma-Aldrich, S2378) or low-density polyethylene (LDPE FT5230, M_w_ of 97,300 g/mol, Borealis, Austria). Each liter of MM (pH 7.2) contained 100 mL 10x M9 Medium (0.56 M Na_2_HPO_4_, 0.29 M KH_2_PO_4_, 85.56 mM NaCl and 93.47 mM NH_4_Cl), 10 mL 100x Trace elements solution (17.11 mM, EDTA, pH 7.5, 3.07 mM FeCl_3_×6H_2_O, 0.62 mM ZnCl_2_, 76.26 µM CuCl_2_×2H_2_O, 42.03 µM CoCl_2_×6H_2_O, 0.16 mM H_3_BO_3_ and 8.08 µM MnCl_2_×4H_2_O), 1 mL 1 M MgSO_4_, 0.3 mL 1 M CaCl_2_, 1 mL 1 mg/mL biotin and 1 mL 1 mg/mL thiamin. A 200 µL aliquot of overnight culture was used to inoculate 20 mL of MM supplemented with the substrate to be tested. These precultures were sub-cultured at least three times on the same individual substrate to allow cells to adapt to the specific single carbon source before inoculating the final cultures used for growth profiling and proteomic analysis. Growth was assessed by measuring the optical density (absorbance) at 600 nm (OD_600_) at regular intervals for 10 days. Growth experiments were conducted in two biological replicates, each with three technical replicates.

### Phylogenomic analysis of *Rhodococcus* sp. ASF-10

For comparative genomic analysis of *Rhodococcus* sp. ASF-10, publicly available genome sequences of the following 21 bacteria were downloaded from NCBI: *Rhodococcus qingshengii* BF1 (GCA_019279115.1), *Rhodococcus qingshengii* CL-05 (GCA_017910955.1), *Rhodococcus qingshengii* TG-1 (GCA_019048925.1), *Rhodococcus qingshengii* JCM15477 (GCA_001646745.1), *Rhodococcus qingshengii* 7B (GCA_015034605.1), *Rhodococcus qingshengii* RL1 (GCA_008306195.1), *Rhodococcus qingshengii* A34 (GCA_031733215.1), *Rhodococcus* sp. C-2 (GCA_027854095.1), *Rhodococcus erythropolis* R138 (GCA_000696675.2), *Rhodococcus globerulus* NBRC14531 (GCA_001894805.1), *Rhodococcus wratislaviensis* NBRC100605 (GCA_000583735.1), *Rhodococcus koreensis* DSM 44498 (GCA_900105905.1), *Rhodococcus jostii* DSM 44719 (GCA_900105375.1), *Rhodococcus opacus* DSM 43205 (GCA_910591545.1) *Rhodococcus opacus* R7 (GCA_000736435.1), *Rhodococcus ruber* C1 (GCA_016804345.1), *Rhodococcus xishaensis* LHW51113 (GCA_004011825.1), *Rhodococcus spelaei* C9-5 (GCA_006704125.1), *Rhodococcus maanshanensis* DSM 44675 (GCA_900109405.1), *Rhodococcoides trifolii* CCM 7905 (GCA_014635345.1), and *Corynebacterium antarcticum* CCM 8835 (GCA_016595285.2). Taxonomic assignment of all genomes was performed using GTDB-Tk v2.4.0 (12) and GTDB database release 220 (13). The phylogenetic tree was generated using the concatenated alignment of 120 ubiquitous single-copy proteins obtained from GTDB-Tk, followed by the tree inference from the multiple sequence alignment with settings ‘--prot_model WAG --gamma’. The tree was rooted employing *Corynebacterium antarcticum* CCM 8835 (GCA_016595285.2) as an outgroup and visualized using R v4.3.3 (14) and the R packages tidyverse (15), ggtree (16), tidytree (17) and ape (18). For *Rhodococcus* sp. ASF-10 and *R. trifolii* CCM 7905, ANI values were calculated using FastANI v1.34 (19). AAI values were determined using FastAAI v0.1.20 (20).

### Comparative genomic and functional trait analysis of *Rhodococcus* genomes

Metabolic complexity across the 21 *Rhodococcus* genomes was assessed using distillR v0.3.0 (21), a R package that transforms raw annotations into genome-inferred functional traits (GIFTs). DistillR incorporates over 300 curated metabolic pathways and modules from the KEGG (22) and MetaCyc (23) databases to quantify microbial metabolic potential based on the relative abundance of pathway-associated genes. For each genome, GIFT-derived metabolic capacity values are averaged to generate a genome-level metric of overall metabolic capacity, hereafter referred to as the Metabolic Capacity Index (MCI). The *C. antarcticum* and *Rhodococcus* genomes were first annotated using DRAM v1.5 (24) to obtain KEGG Orthology (KO) identifiers, which were then mapped to GIFT functional elements followed by MCI calculations. Genomes were categorized into three classes based on total genome size: small (<6 Mb), medium (6-8 Mb), and large (>8 Mb). To test for significant differences in metabolic complexity across genome size categories a one-way analysis of variance (ANOVA) with MCI as the dependent variable and genome size category as the independent variable was performed. Post-hoc pairwise comparisons were conducted using Tukey’s Honestly Significant Difference (HSD) with a confidence level set to 0.95 and family-wise error rate correction applied to control for multiple comparisons. To visualize metabolic functional profiles, we performed t-distributed Stochastic Neighbor Embedding (t-SNE) analysis on the GIFT functional elements matrix using the R package Rtsne v0.15 (25). Perplexity was set to the minimum of 30 or (n-1)/3 to accommodate the sample size. All analyses were performed in R v4.3.3 (14), and results were visualized using ggplot2 v3.5.2 (26), cowplot v1.2.0 (27) and pheatmap v1.0.13 (28).

### Genomic potential for the degradation of synthetic and natural polymers

The genome of *Rhodococcus* sp. ASF-10, along with the genomes of the other 20 *Rhodococcus* spp. used for phylogenomic analysis, was screened for 593 unique gene products associated with the degradation of a selection of synthetic and biodegradable polymers, as well as alkanes. The latter substrate was chosen since the products arising from abiotic degradation of synthetic polymers are identical to or resemble alkanes and other forms of hydrocarbons found in nature (both fossil and renewable). The list of these proteins was compiled from the PlasticDB database (29), the Plastics-Active Enzymes Database PAZy (30), the Hydrocarbon Aerobic Degradation Enzymes and Genes HADEG (31) and published literature (32–34). The target polymers considered in this search included PE as well as derivatives and oxidized derivatives of PE (alkanes and 2-ketones), polystyrene (PS), polyurethane (PUR), polyamide (PA), polyethylene terephthalate (PET), polyvinyl alcohol (PVA), polyethylene glycol (PEG), polycaprolactone (PCL), polylactic acid (PLA), polyhydroxyalkanoates (PHA), polyethylene furanoate (PEF), polybutyl succinate (PBS), poly(butylene succinate-co-adipate) (PBSA), polybutylene adipate terephthalate (PBAT) and natural rubber (NR). The subject database consisted of the predicted proteome of *Rhodococcus* sp. ASF-10, the predicted proteome of the 20 *Rhodococcus* spp. used for genomic analyses as well as the predicted proteome of *C. antarcticum* CCM 8835. The amino acid sequences of the proteins encoded by 593 genes associated with plastic- and hydrocarbon degradation were used as query to search the subject database using Protein-Protein BLAST v2.16.0+ (35) with settings ‘-max_target_seqs 593’ (number of sequences in the query dataset). BlastP hits were filtered for ≥50% identity (pident), ≥75% query coverage (qcovhsp) and e-value of ≤1.0e-5. For each protein, the best remaining hits were retained. Results were visualized using ggplot2 v3.5.2 (26) in R v4.3.3 (14).

### Assessment of LMWPE degradation by *Rhodococcus* sp. ASF-10

Three analytical techniques were used to gain insight into the effects of *Rhodococcus* sp. ASF-10’s growth on LMWPE structure and composition. To prepare samples for analysis, *Rhodococcus* sp. ASF-10 was cultured in triplicates in MM supplemented with 30 mg/mL LMWPE for 7 days at RT (referred as *Rh*LMWPE). As controls, cell-free MM was incubated with 30 mg/mL LMWPE (referred as cfLMWPE). Untreated LMWPE, (referred as uLMWPE), that was not incubated in MM, was also included in the analysis. Following incubation, LMWPE powder was separated from the MM by filtration with Whatman 20 µm cellulose filters (Cytiva, USA, Cat. No: WHA10331554). *Rh*LMWPE was washed in three consecutive steps to remove biofilm. In the first step, 500 µL dH_2_O was added, and the LMWPE was kept in a ThermoMixer (Eppendorf, Germany) at 1500 rpm for 30 min at RT. This was followed by an ultrasonication step in a Bransonic 3510-DHT Ultrasonic Bath (Emerson, USA) for 10 min, before removing the dH_2_O. The cleaning was then repeated using 500 µL of a 1 M NaOH solution, followed by washing with 500 µL of 70% ethanol, and the washed LMWPE particles were dried in a Concentrator plus (Eppendorf, Germany) at 45°C for 1 h. The control samples cfLMWPE and uLMWPE were subjected to identical treatment.

Fourier-transform infrared spectroscopy (FT-IR) was first performed using a Spectrum Two FT-IR spectrometer (PerkinElmer, Waltham, MA, USA) to detect potential variations in functional groups along the polymer backbone. The signals were obtained with 4 cm^-1^ spectral resolution (8 consecutive readings per measurement) in the 4000-550 cm^-1^ range using Spectrum IR software (PerkinElmer, Waltham, MA, USA).

Size exclusion chromatography (SEC) was performed using a GPC-IR5 system (Polymer Char, Valencia, Spain) to determine changes in the size of the polymer chains. Approximately 4 mg of each sample was dissolved in 8 mL of 1,2,4-trichlorobenzene at 160 °C for 3 hours. A 200 µL aliquot was injected into the size exclusion chromatography SEC system equipped with four PLgel 20 µm MIXED-A columns (Agilent Technologies, Santa Clara, CA, USA). Analysis was conducted at 150 °C using 1,2,4-trichlorobenzene as the mobile phase (flow rate: 1 mL/min) and a high-sensitivity infrared detector. Calibration was performed using polystyrene standards with narrow molecular mass distributions and peak molecular weights (M_peak_) ranging from 1,140 to 7,500,000 g/mol. The weight average molecular weight (M_w_), number average molecular mass (M_n_), and molar mass distribution of the materials were determined.

Finally, changes in the amounts of low-molecular-weight compounds (alkanes and ketones) in the samples were determined using gas chromatography-mass spectrometry (GC-MS). For extraction, approximately 100 mg of sample was mixed with 3 mL ethyl acetate in a 4 mL glass vial, sealed with a PTFE/Silicone septum, and heated at 95°C for 1.5 h. The extract was filtered (0.2 µm Teflon syringe filter) and analyzed using an Agilent 6890N GC with a 5973 MS detector and GERSTEL MPS2 autosampler. Compounds were separated on a Zebron ZB-5MSPlus column (30 m × 250 µm, 0.25 µm film). Oven temperature was ramped from 60 °C to 300 °C at 10 °C/min over 45 min. Helium (grade 6.0) was used as carrier gas (3 mL/min). Injections were splitless at 250 °C; ion source temperature was 230 °C; and ionization was by EI at 70 eV. Spectra were acquired from m/z 33–720. Each sample was run in triplicate, and compounds were identified using the Wiley 11th/NIST 2017 spectra library. Compound quantification was carried out using the reference compounds BHT (butylated hydroxytoluene) and Tinuvin 120 (2’,4’-Di-tert-butylphenyl 3,5-di-tert-butyl-4-hydroxybenzoate), which give differences in responses between lower and higher molecular weight components. Specifically, the response of BHT falls in the region of lower molecular weight alkanes and ketones (up to C16), while Tinuvin 120 falls in the range of higher molecular weight alkanes and ketones. The analyzed LMWPE samples were confirmed to contain no compounds chemically similar to Tinuvin 120 or BHT.

### Analysis of *Rhodococcus* sp. ASF-10’s proteome in response to oxidized LMWPE

*Rhodococcus* sp. ASF-10 was grown in triplicate on MM supplemented with either 2% (w/v) sodium succinate or 3% (w/v) LMWPE, respectively, as a sole carbon source. Samples (20 mL) were harvested at the mid-exponential growth phase. Planktonic cells were collected from the liquid cultures by centrifugation (4,500 × g, 10 min, 4°C), resuspended in 50mM Tris-HCl pH 7.5, 100 mM NaCl, 0.1% (v/v) Triton X-100, 200mM NaCl, 1 mM dithiothreitol (DTT) and disrupted by bead-beating using three 60 s cycles at 6.5 m/s with a FastPrep24 (MP Biomedicals, CA). Cell debris was removed by centrifugation (16,000 × g, 10 min, 4°C), and proteins were precipitated with ice-cold trichloroacetic acid (TCA), final concentration of 10% (v/v), incubated on ice overnight, centrifuged (15,000 × g, 15 min, 4°C) to pellet the precipitated proteins and washed with 300 μL ice-cold 0.01M HCl in 90% acetone. After removal of the supernatant, the protein pellets were air-dried and resuspended in a buffer containing 5% SDS and 50 mM triethylammonium bicarbonate (pH 8.5). Protein digestion was performed using S-Trap Mini Columns (Protifi, Fairport, NY, USA) according to the manufacturer’s instructions, with 480 mM DTT for reduction and 1 M iodoacetamide (IAA) for alkylation. The resulting peptides were analyzed on a nanoLC-MS/MS system, consisting of a nano UHPLC (nanoElute 2, Bruker Daltonics Inc., Bremen, Germany) coupled to a trapped ion mobility spectrometry/quadrupole time of flight mass spectrometer (timsTOF Pro, Bruker Daltonics Inc., Bremen, Germany). Peptides were separated on an Aurora C18 reverse-phase (1.6 µm, 120 Å) 25 cm x 75 µm analytical column with an integrated emitter (IonOpticks, Melbourne, Australia). Mass spectral data were acquired using DataAnalysis v6.1.

MS raw files were processed using the FragPipe v21.1 (with MSFragger v4.0, IonQuant v1.10.12, Philosopher v5.1.0, DIA-NN v1.8.2 beta 8) (36–39) for protein identification and label-free quantification (LFQ). Protein identification was performed by searching MS and MS/MS spectra against the complete proteome of *Rhodococcus* sp. ASF-10 (5,538 sequences), supplemented with common contaminants (e.g., keratins, trypsin, and bovine serum albumin, 118 sequences). To estimate false discovery rates (FDRs), a decoy database consisting of reversed sequences of all protein entries was included (5,656 sequences). The resulting protein library consisted of 11,312 proteins. Trypsin was specified as the proteolytic enzyme, allowing one missed cleavage. Variable modifications included N-terminal acetylation, methionine oxidation, deamidation of asparagine and glutamine residues, and pyro-glutamate formation at N-terminal glutamines, while carbamidomethylation of cysteines was set as a fixed modification. Protein identifications were filtered to achieve a 1% FDR. A protein was considered ‘present’ if detected in at least two of the three biological replicates for at least one substrate. Missing values were imputed from a normal distribution (width: 0.3; downshift: 1.8 standard deviations from the original distribution) using the “total matrix mode”. Differential abundance analysis was conducted with Rstatix v0.7.2 (40) using an unpaired two-tailed student’s t-test with a permutation-based FDR threshold set at 0.05. Figures were generated using R v4.3.3 (14) with the packages ggplot2 v3.5.1 (26) and cowplot v1.1.3 (27).

### Weighted gene co-expression network analysis

A protein network was constructed using weighted gene co-expression network analysis (WGCNA) as an unsupervised method. Briefly, log₂-transformed proteomic data, including imputed missing values, was used to construct the network and identify protein modules using the R package WGCNA v1.73 (41), which employs the random matrix theory approach. The soft-thresholding (β) was selected using the “pickSoftThreshold” function to approximate scale-free topology (β = 20, R2 = 0.864, mean connectivity = 321). An adjacency matrix was computed with networkType = “signed”, transformed into a topological overlap matrix (TOM), and converted to a dissimilarity matrix. Hierarchical clustering of proteins based on TOM dissimilarity was conducted using average linkage, and modules were defined with the cutreeDynamic function with parameters minClusterSize = 30 and deepSplit = 2. Subsequently, closely related modules were merged using the function mergeCloseModules at a cut height of 0.25. Module eigengenes were calculated, clustered, and correlated with growing substrate (sodium succinate or oxidized LMWPE) to assess module-substrate associations. Module significance was estimated by t-test and adjusted *P* values (FDR).

Relevant pathways for this study, such as alkane and ketone degradation, fatty acid transporter systems, fatty acid metabolism, biofilm formation and surfactant production, were identified for proteins annotated with DRAM that had assigned a KEGG Ontology (KO) identifier using the KEGGREST R package (42). The identified proteins in the pathways listed above were intersected with the proteins in the module positively associated with LMWPE using the R package UpSetR (43).

## Results and discussion

Among the bacterial genera implicated in hydrocarbon biodegradation, *Rhodococcus* stands out alongside *Bacillus* and *Pseudomonas* as one of the key contributors to this process (44, 45). *Rhodococcus* spp. have been isolated from a variety of ecosystems, including soil contaminated with hydrocarbons or plastics, waste sites with naturally weathered plastic debris, seawater and industrial wastewater (33, 44, 46, 47). Comparative genomic analysis across the *Rhodococcus* genus has identified a diverse set of oxidative systems, including multicopper oxidases, alkane monooxygenases, cytochrome P450 hydroxylases, para-nitrobenzylesterases, and carboxylesterases that have been postulated as compatible with the ability of depolymerize plastics with C–C backbones, those containing heteroatoms in the main chain, and various polyesters (48).

Bacteria of the genus *Rhodococcus* are commonly present as part of the gut microbiota of salmon across all developmental stages, from fry to adults (49). In an effort to identify and characterize salmon gut bacteria interacting with microplastics components that are common in aquaculture, such as PE and degradation products arising from abiotic oxidation of the polymer, we found that *Rhodococcus* sp. ASF-10, isolated from the gut of Atlantic salmon (11), displayed the ability to grow on a LMWPE (**Figure 1**). This compound was previously found to be oxidized (50), and can be considered a model substrate for abiotically degraded PE.

**Figure 1.**
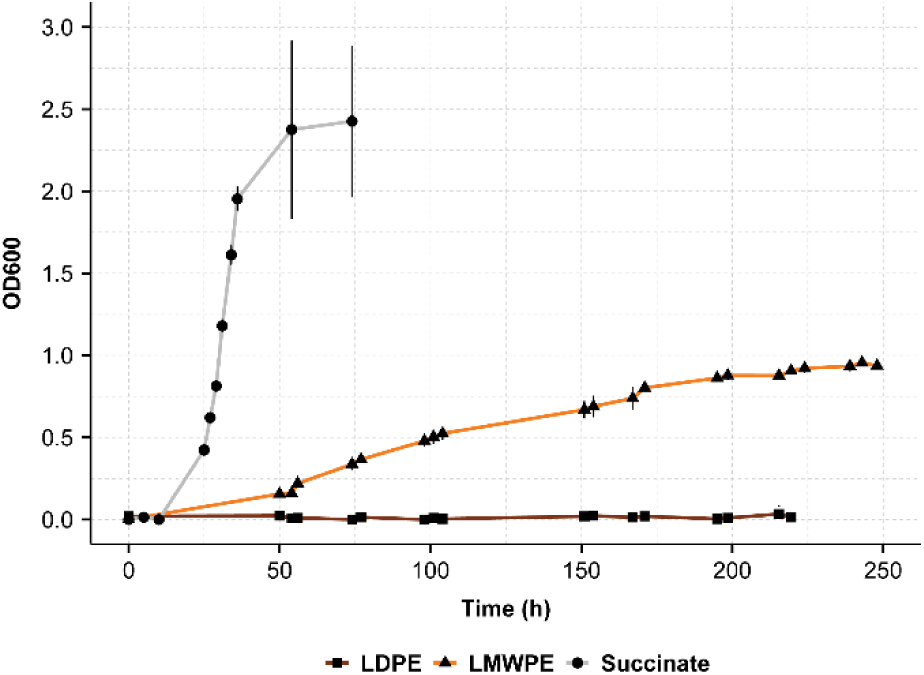
Growth curves for *Rhodococcus* sp. ASF-10. Growth was measured as the optical density at 600 nm (OD600). The values represent the averages of two biological replicates with technical triplicates. Abbreviations: LDPE, low density polyethylene; LMWPE, low molecular weight polyethylene.

### Comparative genomic analyses and functional profiling of *Rhodococcus* isolates reveal conserved metabolic traits

To explore the metabolic potential of *Rhodococcus sp.* ASF-10 for possible utilization of plastics, hydrocarbons and their oxidized derivatives, we first determined the phylogenomic relationship of our isolate and compared its genomic features with those of previously described *Rhodococcus* isolates. Indeed, many members of the genus *Rhodocuccus* have been associated with the degradation of hydrocarbons, including alkanes, and PE. These include *R. qingshengii*, *R. erythropolis*, *R. opacus*, *R. aetherivorans* and *R. jostii* (51) reported to degrade alkanes, and some *Rhodococcus* are hypothesized to be able to act upon and depolymerize LDPE, including *R. opacus*, *R. ruber*, *R. qingshengii* and *R. equi* (44, 46, 52, 53).

To establish the evolutionary relationship between *Rhodococcus* sp. ASF-10 and 20 other members of the genus isolated from a diverse set of environments, a phylogenetic tree based on 120 conserved, single-copy markers shared across the species was constructed, resolving six distinct clades (**Figure 2**). *Rhodococcus* sp. ASF-10 exhibited the closest evolutionary relationship to *R. trifolii* CCM 7905, a strain originally isolated from the leaf surface of a plant (54) (clade 3 in **Figure 2**). Following comparisons of the two genome sequences, an ANI (Average Nucleotide Identity) value of 78.92 and an AAI (Average Amino Acid Identity) value of 65.03 indicated that the two species were distantly related. Among the other *Rhodococcus* spp., 10 isolates have previously been associated with either alkane (55), crude oil (56) or PE degradation (33, 47, 52). These include all the *R. qingshengii* strains, except one (*R. qingshengii* JCM15477), and indeed, they all showed a close evolutionary relationship in our analysis (clade 1 in **Figure 2**). Clade 2 and 4 in the phylogenomic tree consisted of soil and water isolates of which only *R. opacus* R7 has been reported to be associated with PE utilization (33). Clade 6 included *R. ruber* C1, an isolate known to degrade long-chain alkanes (55). The genomes of the *Rhodococcus* spp. showed variations in genome size, ranging from *R. xishaensis* LHW51113 with a genome of 3.7 Mb to *R. wratislaviensis* NBRC100605 with a genome of 10.4 Mb. Strains within the same clade generally had similar genome sizes, with genomes in clade 2 being between 8.2-10.4 Mb (large size genomes), in clade 1 between 6.3-7.8 Mb (medium size genomes), and the remaining strains have genomes of <6 Mb (small size genomes). *Rhodococcus sp.* ASF-10 belongs to the group of isolates with small genomes.

**Figure 2.**
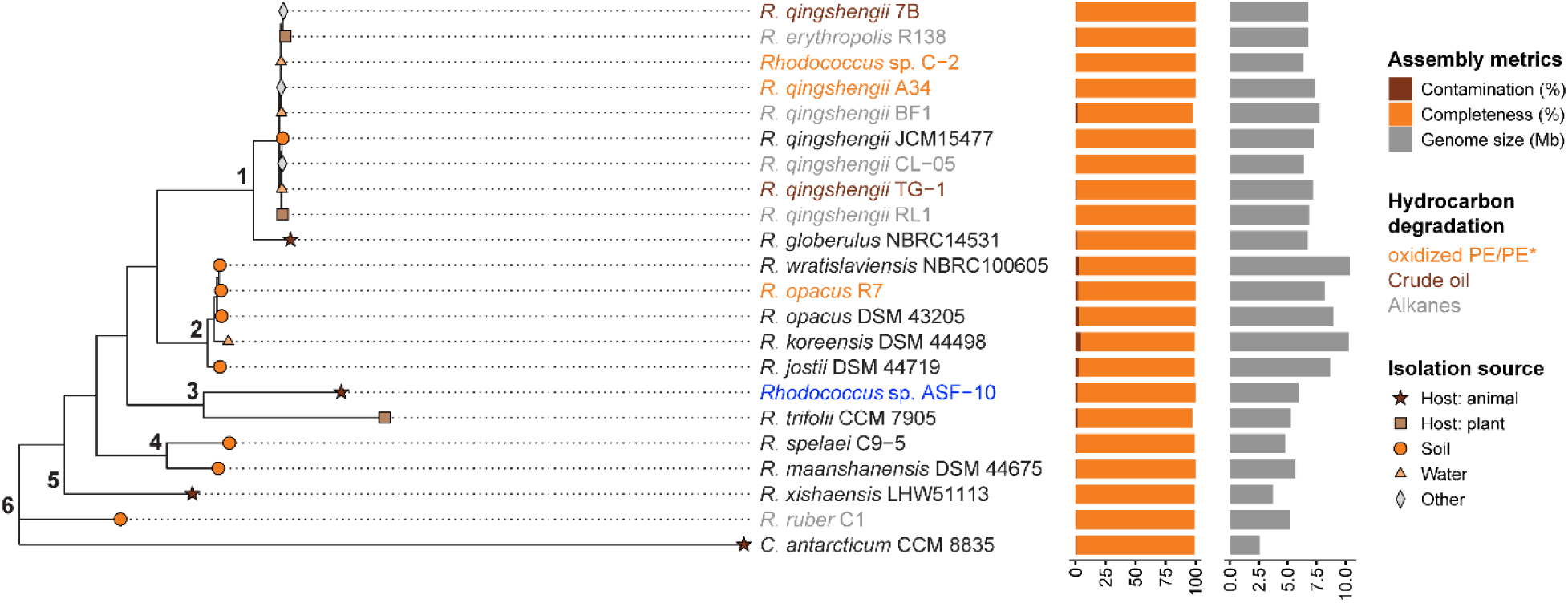
Phylogenomic tree displaying the evolutionary relationships of *Rhodococcus* sp. ASF-10 from this study (highlighted in blue) and 20 publicly available *Rhodococcus* genomes. The genome of *Corynebacterium antarcticum* CCM 8835 (GCA_016595285.2) was used as the outgroup. Numbers (1–6) denotes the six clades represented in the tree. Different coloring of the strain names depicts their known ability to degrade or interact with different hydrocarbons (alkanes, oil and polyethylene). The horizontal bar plots displayed next to each genome depict genome quality (orange bars for completeness and brown bars for contamination), and genome size (grey bars). The * in “oxidized PE/PE” indicates that the substrate used in the studies that characterize this ability in the corresponding bacteria may have contained low-molecular weight PE-derivatives. Abbreviations: PE. polyethylene.

To compare specific metabolic capacities encoded by *Rhodococcus sp.* ASF-10 and the 20 selected *Rhodococcus* genomes, we computed the functional potential of each genome based on annotated KEGG Orthology IDs (KO IDs), that were transformed into genome-inferred functional traits (GIFTs). Genome ordination based on GIFTs values revealed clustering patterns of the *Rhodococcus* species (**Figure 3A-B**). Genomes in clade 2 (*R. opacus* ssp., *R. wratislaviensis* and *R. koreensis*) formed a distinct cluster. In addition, genomes in clade 1 (*R. qingshengii* ssp.) also grouped closely together. No clear clustering pattern according to the source of isolation was observed. This is consistent with the ecological versatility of *Rhodococcus*, which inhabits diverse environments ranging from soil and water to host-associated niches (57). Species clustering aligned with the clades that were assigned in the phylogenomic analysis (**Figure 2**), indicating that phylogenomic relationship largely determines the genome grouping and metabolic traits (**Figure 3A**). This relationship was also evident when looking at the metabolic capacity indices (MCIs) of each strain (**Figure 3B**). A significant positive relationship was observed between MCI values and genome size. By grouping the genomes by sizes, an analysis of variance (ANOVA) and post-hoc analysis (Tukey HSD) confirmed that large and medium genomes tend to exhibit higher MCI values than small genomes. Comparisons of large versus small genomes (*P* < 0.01) as well as medium versus small genomes (*P* < 0.01) revealed a highly significant correlation between genome size and metabolic complexity. Genomes with sizes ranging from 6-8 Mb and size >8 Mb showed very similar MCIs, with average MCIs of 0.27 ± 0.006 and 0.27 ± 0.003, respectively. Indeed, no significant correlation between large and medium genome sizes and their MCIs was observed for these two groups. Small genomes showed a significantly lower metabolic complexity (average MCI 0.24 ± 0.02) compared to large and medium size genomes. The lack of a significant difference in MCI values between medium and large genomes suggests a threshold (∼6 Mb in genome size) beyond which further increases in genome size do not confer additional metabolic capacity. This may suggest that core metabolic functions are established by 6 Mb, and that additional genome content beyond 6 Mb may be redundant genes, mobile elements, or adaptive traits that enable survival across varied environments rather than increased metabolic complexity.

**Figure 3.**
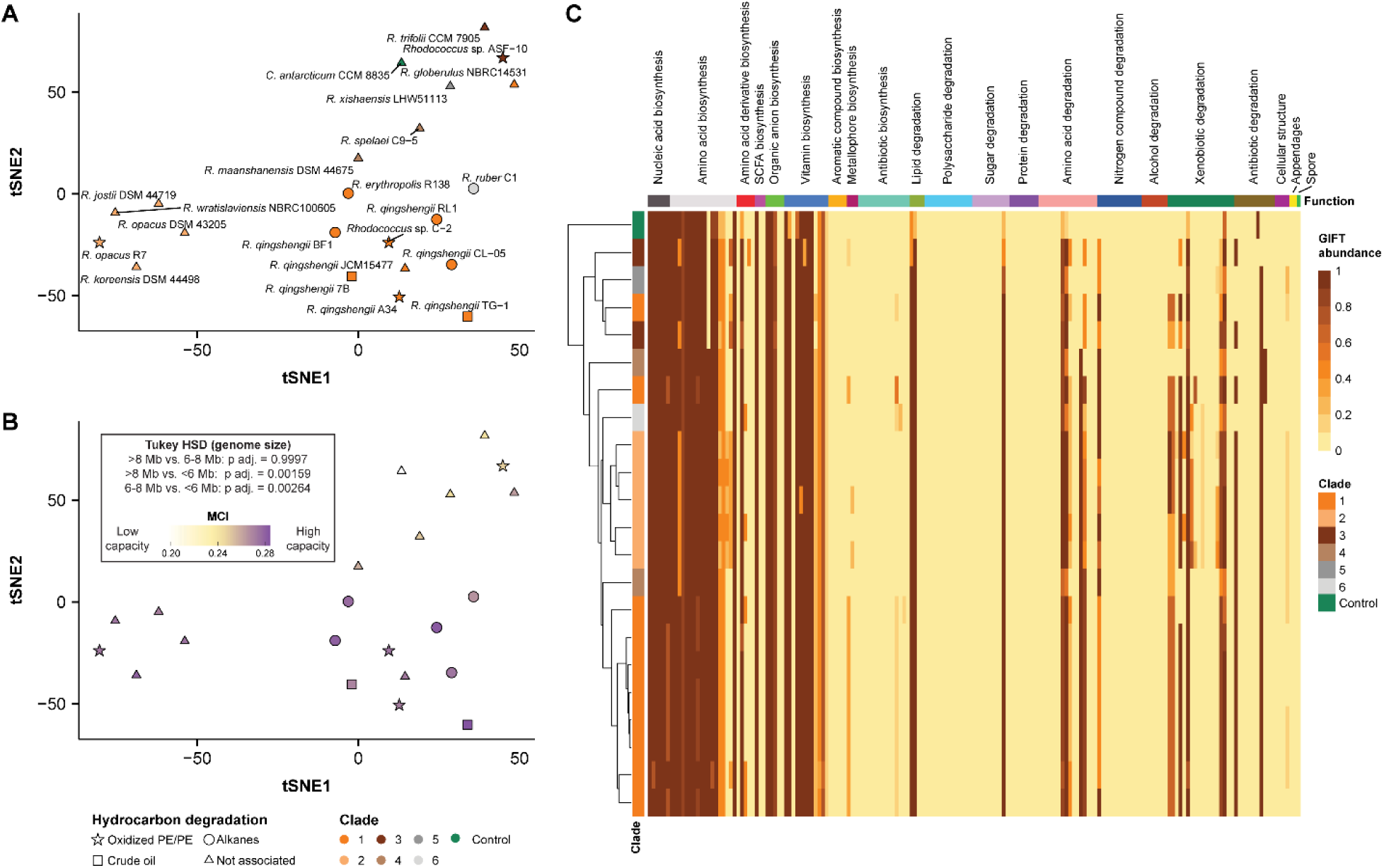
Functional ordination of *Rhodococcus* genomes based on GIFTs and comparison of metabolic trait distributions. (**A**) Ordination of the *Rhodococcus* genomes based on GIFT values using a t-Distributed Stochastic Neighbor Embedding (t-SNE) analysis. Genomes are colored according to the clade assignment from the phylogenomic analysis. (**B**) Same ordination as in (**A**), but genomes are colored by their metabolic capacity index (MCI). Lowest complexity genomes are found in the bottom left, and highest complexity genomes are found in the top right. An ANOVA test and post-hoc test (Tukey) was performed to analyze the differences in MCI across strains with different genome sizes. (**C**) Heatmap of GIFT abundance associated with a particular metabolic functions (columns) in each *Rhodococcus* genomes analysed in this study. Genomes are clustered based on GIFT abundance (left dendogram). Abbreviations: PE. polyethylene.

Next, we compared the biosynthetic and degradative capacities for specific biomolecules across the *Rhodococcus* strains (**Figure 3C**). Despite no drastic differences being observed across most pathways, certain metabolic functions varied among clades. Specifically, clade 2 genomes include pathways for amino acid biosynthesis and xenobiotic degradation that are not found in the genomes belonging to the other clades (**Figure 3C**). Clade 3, that includes *Rhodococcus* ASF-10 (small genome size), was characterized by the lack of pathways for production of glutamate, gamma-aminobutyric acid (GABA), and vitamin B3 (niacin). Otherwise, the core functions of all *Rhodococcus* genomes are generally the same. They all have the complete pathway for degradation of lipids, specifically fatty acid degradation. Importantly, this pathway includes enzymes for utilization of alkanes, which seems to be a core function for *Rhodococcus* spp.

Overall, our results underscore that phylogenomic relationships largely govern metabolic trait patterns in *Rhodococcus* genomes, with genome size directly correlated with metabolic complexity up to a threshold of approximately 6 Mb. Remarkably, degradation of fatty acids, and the biosynthesis of multiple nucleic acids (inosinic acid, uracil, cytosine and adenine) and amino acids (serine, threonine, valine, leucine, proline, tyrosine), as well as the vitamin B2 (riboflavin) are conserved among the *Rhodococcus* spp., including *Rhodococcus* ASF-10. Other pathways, which are mostly conserved across the genomes, include the biosynthesis of the short-chain fatty acid acetate, organic acids (succinate, fumarate and citrate), several B vitamins and most amino acids, as well as the degradation of xenobiotics such as toluene, benzoate, catechol and trans-cinnamate.

### Genomic potential for degradation of hydrocarbons in *Rhodococcus* spp

To assess the *Rhodococcus* strains’ potential to utilize hydrocarbons, we generated a reference database that includes 593 unique proteins associated with the utilization of diverse synthetic and biodegradable polymers, as well as alkanes. The list of these proteins was compiled from the PlasticDB database (29), the Plastics-Active Enzymes Database PAZy (30), the Hydrocarbon Aerobic Degradation Enzymes and Genes HADEG (31) and published literature (32–34). BLAST searches indicated that all *Rhodococcus* genomes encoded proteins with ≥50% sequence identity to the proteins in the reference databases, while, as expected, no hits were obtained for *C. antarcticum* (**Figure 4** and **Table S1**). The capacity to metabolize or interact with alkanes appeared to be broadly conserved across the *Rhodococcus* genus. (**Figure 4**). In general, closely related strains shared similar potential for alkane degradation. For example, all the *R. qingshengii* strains, along with *R. globerulus* NBRC14531 in clade 1, all have hits in their gene encoding alkane 1-monooxygenase (AlkB), long-chain alkane monooxygenase (LadA) and a FAD-binding monooxygenase associated with long-chain alkane degradation (AlmA), as well as the electron-transfer protein rubredoxin (RubA). These enzymes are known to be involved in the initial terminal/biterminal hydroxylation of alkanes (58). Clade 2, consisting of *R. wratislaviensis* NBRC100605, *R. opacus* R7*, R. opacus* DSM 43205*, R. koreensis* DSM 44498 and *R. jostii* DSM 44719 (**Figure 2**) have genes coding for enzymes for both subterminal and terminal/biterminal oxidation of alkanes (**Figure 4**). In *Rhodococcus* sp. ASF-10, we detected 12 genes coding for proteins associated with alkane degradation, particularly those involved in terminal/biterminal oxidation of alkanes and 4 with plastic degradation, specifically PE, PUR and PA. Overall, the species within the same clade generally have genes encoding for a main degradation mechanism, with clade 1, 3, 4 and 5 mainly using terminal/biterminal oxidation and clades 2 and 6 using subterminal oxidation.

**Figure 4.**
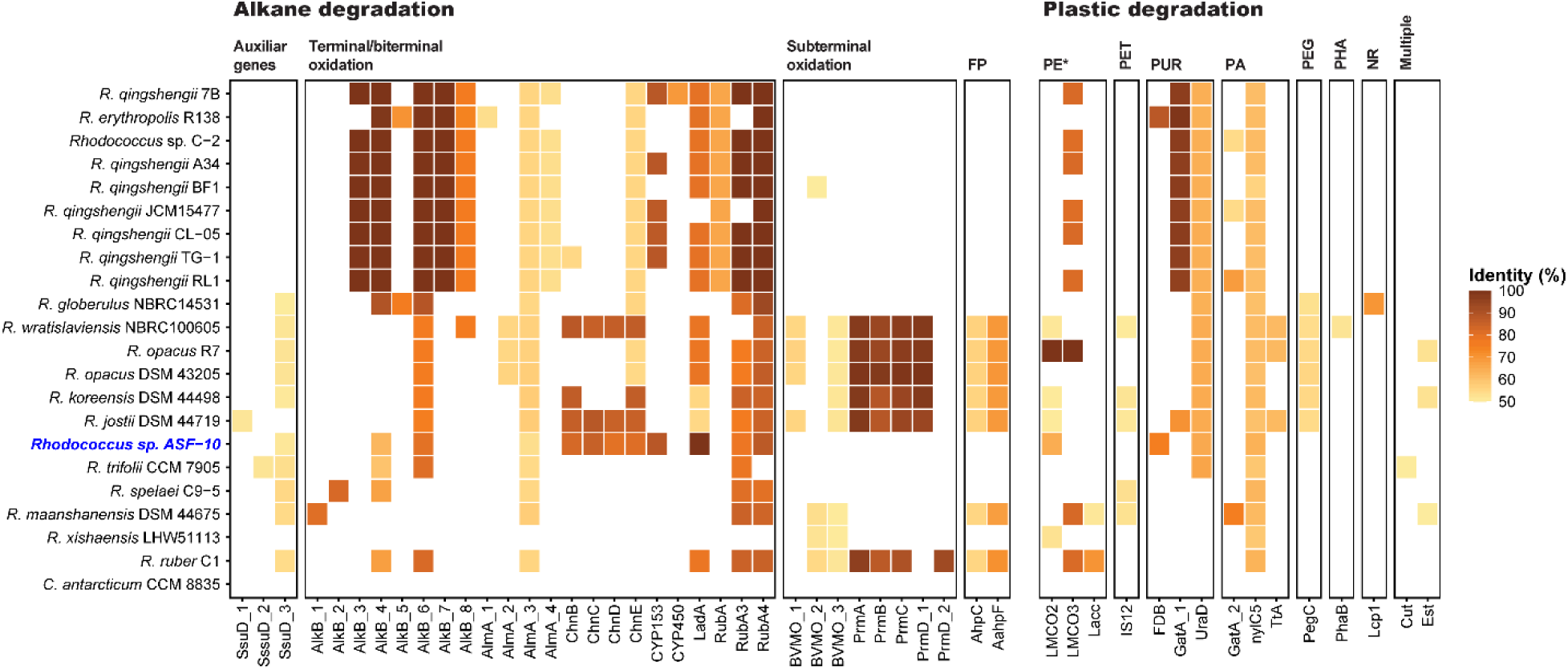
Detection of enzymes for alkane degradation, fossil-based plastics and biodegradable plastics in the predicted proteome of *Rhodococcus* sp. ASF-10 and 20 publicly available *Rhodococcus* genomes. The predicted proteome of *Corynebacterium antarcticum* CCM 8835 was included as a control in the bioinformatic analysis, as the genome of this bacterium is known to lack the targeted genes. Hits represent best match in each proteome. The * in “PE” indicates that the substrate used in the studies that characterize this ability in the corresponding bacteria to may have contained low-molecular weight PE-derivatives. Abbreviations: polyethylene terephthalate (PET), polyurethane (PUR), polyamide (PA), polyethylene glycol (PEG), polyhydroxyalkanoates (PHA), natural rubber (NR). The panel “multiple” contains hydrolases which act on multiple plastics (the cutinase Cut on PCL/PPA and the lipase Est on PHBV/PHO/PLA/PCL/PES). BlastP hits and metadata from PAZy, HADEG and PlasticDB are provided in **Table S1**.

Though 489 of the 593 entries in our database were proteins related to the degradation of synthetic polymers and biopolymers, only 16 of the gene encoding plastic-associated enzymes were detected in the *Rhodococcus* genomes (**Figure 4** and **Table S1**). Some of the *Rhodococcus* spp., including the *Rhodococcus* sp. ASF-10 from this study, had hits for two laccase-like multicopper oxidases (LMCO) LMCO2 and LMCO3, which were previously associated with the degradation of PE (33, 53). Another copper-binding enzyme, a laccase (Lacc) (46), was detected in the strain *R. ruber* C1 and *R. maanshanensis* DSM 44675. Although these enzymes have been linked to the degradation of PE, Lacc has not yet been biochemically confirmed as a PE-degrading enzyme. Notably, the LMCO2 and LMCO3 were characterized using the same LMWPE material used in this study (Sigma-Aldrich, 427772), which has been shown to contain PE-derivatives (alkanes) and oxidized PE-derivatives (ketones), in addition to the polymeric component of LMWPE (50). This opens the possibility that the two LMCOs could be active only on the alkanes and ketones present in the substrate mixture. All *Rhodococcus* strains previously associated with PE degradation possess putative homologs for one or two of these enzymes (**Figure 4**). However, in the absence of experimental evidence confirming actual degradation of the polymeric component in LMWPE, the presence of the genes coding for the laccases and LMCOs does not confirm a true capacity for PE degradation.

Next, we explored the genomic potential of the *Rhodococcus* spp. to interact with other synthetic and biodegradable polymers (**Figure 4**). The genomes of *R. wratislaviensis* NBRC100605, *R. koreensis* DSM 44498, *R. jostii* DSM 44719, *R. spelaei* C9-5 and *R. maanshanensis* DSM 44675 includes genes encoding putative enzymes for PET degradation (cutinase IS12, (59)). For PUR utilization, *R. erythropolis* R139 and *Rhodococcus* sp. ASF-10 displayed amidases (FDB, (60)), while the putative ability to utilize PUR through the urethanase GatA_1 (60) was detected among all the *R. quingshengii* ssp. and *R. erythropolis* R138. Seventeen out the 20 *Rhocodoccus* spp. examined possessed a protein similar to the esterase UraD, previously associated with PUR degradation (60). Regarding the utilization of PA, while a protein highly similar to the amidase nylC5 for nylon decomposition (61) was detected in all the *Rhocodoccus* spp. GatA_2 for utilization of nylon-6 (62) was detected only in the proteome of *Rhodococcus* sp. C-2, *R. quingshengii* JCM15477 and *R. quingshengii* RL1. Finally, the genome of *R. jostii* DSM 44719, *R. opacus* R7 and *R. wratislaviensis* NBRC100605 included a gene coding for a protein similar to the amidase TtA, previously shown to be involved in nylon-6 degradation (62). The potential ability to interact with PHA through the poly(3-hydroxyalkanoate) depolymerase PhaB (63) was detected only in *R. wratislaviensis* NBRC100605, while *R. globerulus* NBRC14531 was the only genome encoding a protein similar to the rubber oxidase Lcp1 for NR degradation (64). Genes coding for enzymes for PEG degradation were identified in the genomes of *R. globerulus* NBRC14531, *R. wratislaviensis* NBRC100605, *R. opacus* ssp., *R. koreensis* DSM 43205 and *R. jostii* DSM 44719.

Overall, our results show that most members of the *Rhodococcus* genus display genomic potential for utilization of alkanes, diverse plastics and natural polymers such as PE, PET, PUR, PA, PEG, PHA and NR, with *Rhodococcus* sp. ASF-10 exhibiting potential for the utilization of alkanes, PE, PUR and nylon.

### Characterization of LMWPE shows that *Rhodococcus* sp. ASF-10 can only utilize alkanes and oxidized derivatives

To determine whether *Rhodoccus* sp. ASF-10 can utilize pristine LDPE or a LMWPE with oxidized derivatives. the isolate was cultured in a MM containing either of these substrates as a sole carbon source. *Rhodoccus* sp. ASF-10 grew well on the oxidized LMWPE, reaching an OD600 of 0.9, whereas no growth was observed on LDPE (**Figure 1**). Substrate utilization by the isolate was examined by characterizing the LMWPE substrate before and after bacterial growth (**Figure 5** and **Table S2**). Firstly, we employed FT-IR spectroscopy that is a commonly used method to determine changes on the surface of the analyzed material, where the formation of carbonyl groups can be detected through the appearance of a peak at ∼1750 cm^-1^. While extensively used to characterize PE post-bacterial growth, FT-IR results are an indirect way to determine surface changes, and should be interpreted with caution, as the existence of oxidized compounds such as ketones and aldehydes in the material, or residual protein or biofilm arising following bacterial growth can result in a signal (65, 66). To circumvent this issue, a washing protocol that included ethanol and sodium hydroxide in combination with sonication was employed. LMWPE was confirmed to contain carbonyl groups (50), which were detected in the untreated material, as well as in the LMWPE incubated in MM with and without *Rhodococcus* sp. ASF-10 as indicated by the presence of peaks at ∼1750 cm^-1^ (**Figure 5A**). No difference was observed between the experimental samples (*Rh*LMWPE). the cell-free samples (cfLMWPE) or untreated controls (uLMWPE), as no additional peaks appeared post-incubation. Another limitation of FT-IR is that it can only detect changes in the outer layers of LMWPE and may not fully capture chemical changes in the deeper layers or reductions in the high-molecular-weight fraction indicative of PE degradation. To address this, the LMWPE material before and after growth with *Rhodococcus* sp. ASF-10 was also analyzed using SEC. The superimposition of the SEC chromatograms of the *Rh*LMWPE and cfLMWPE samples in the high molecular weight region indicated that no degradation occurred in the polymeric fraction of the material, while differences were observed in the low molecular weight region (LogM 2-2.5; **Figure 5B**). To identify the compounds being utilized by *Rhodococcus* sp. ASF-10 during growth, GC-MS analysis was carried out on the LMWPE substrate. After growth with *Rhodococcus* sp. ASF-10 there was a significant reduction (*P* < 0.05) in the amount of 2-ketones and alkanes, particularly of the shorter chain lengths (**Figure 5C**). Overall, *Rhodococcus* sp. ASF-10 was shown to utilize 2-ketones up to 28 carbons and alkanes up to 29 carbons in length.

**Figure 5.**
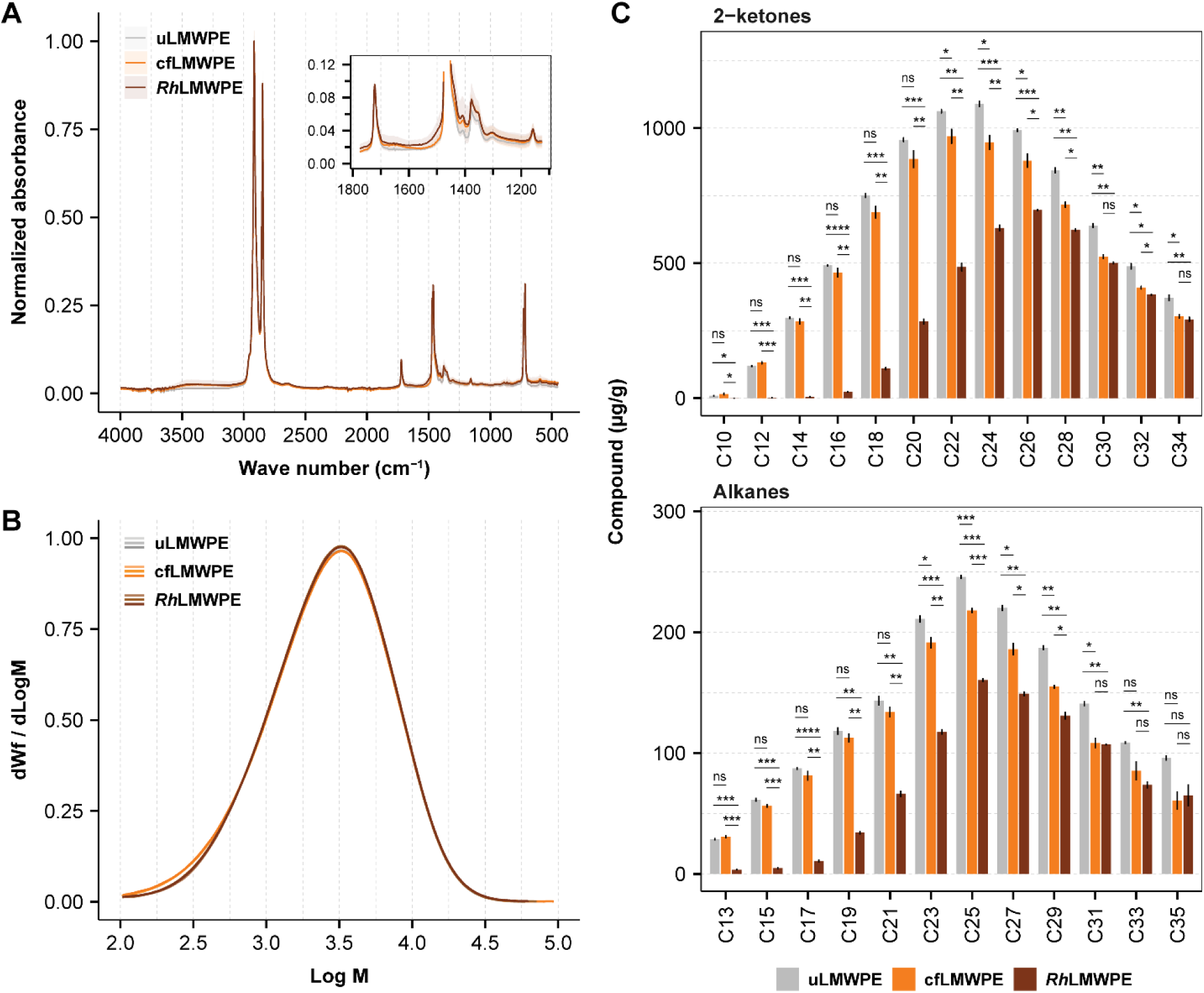
Analyses of LMWPE post-incubation with *Rhodococcus* sp. ASF-10. The isolate was grown with 30 mg/mL on LMWPE in MM (*Rh*LMWPE). Controls consisting of LMWPE incubated in cell-free MM (cfLMWPE) and untreated LMWPE (uLMWPE) were included. **(A)** FT-IR spectra show the mean normalized absorbance of the samples, with the standard deviation included as shaded areas. **(B)** SEC chromatograms of samples. Note that the chromatograms are superimposing in the high molar mass regions, indicating no degradation of the polymeric fraction of the LMWPE substrate, as the distribution would shift to the left. **(C)** GC-MS result of LMWPE samples showing the amount of each compound detected in the LMWPE ethyl acetate extracts. Data are averages ± standard deviations (error bars) of three biological replicates, and significance levels are displayed for pairwise t-test (not significant (ns), p ≤ 0.05 (*), p ≤ 0.01 (**), p ≤ 0.001 (***) and p ≤ 0.0001 (****)). Experiments were conducted in triplicates. Source data for FT-IR, SEC and GC-MS can be found in **Table S2**.

### *Rhodococcus* sp. ASF-10 metabolizes LMWPE derivatives through alkane and fatty acid utilization pathways

Although *Rhodococcus* sp. ASF-10 was unable to utilize the higher molecular weight fraction of LMWPE, it could nonetheless interact with PE-derived compounds occurring in the aquatic environment. To investigate the cellular response under conditions resembling oxidized PE, we carried out a proteomic study using oxidized LMWPE as a model substrate. To identify enzymes involved in the degradation of the LMWPE components, we compared the proteome of *Rhodococcus* sp. ASF-10 grown on this substrate and a sodium succinate reference. We focused on enzymes which have been proposed in previous studies with *Rhodococcus* isolates as being able to depolymerize PE, which belong to different oxidoreductase families, such as alkane hydroxylases and laccases (46, 53, 67). Proteins involved in both terminal and subterminal alkane oxidation pathways were markedly more abundant in the LMWPE-derived proteome than in the succinate-derived proteome. (**Figure 6**). These proteins include an alkane 1-monooxygenase AlkB (AlkB1,2_1_4800), multiple putative cytochrome P450s (CYP_1_2144, CYP_1_3022 and CYP_1_3770), a cytochrome P450 alkane hydroxylase (CYP153_1_3424), as well as several alcohol dehydrogenases and aldehyde dehydrogenases (**Figure 6**). Alkanes are transported into the cell during the first step of the alkane degradation pathway, where AlkB hydroxylates the alkane as the compound is translocated through the cell membrane (68). An additional monooxygenase which can catalyze degradation of long-chain alkanes (AlmA) was detected in the LMWPE proteome (AlmA_1_3897, log_2_ fold-change = 1.29, *P* = 0.07, **Table S3A**), indicating its possible involvement in the processing of long-chain alkanes by *Rhodococcus* sp. ASF-10.

**Figure 6.**
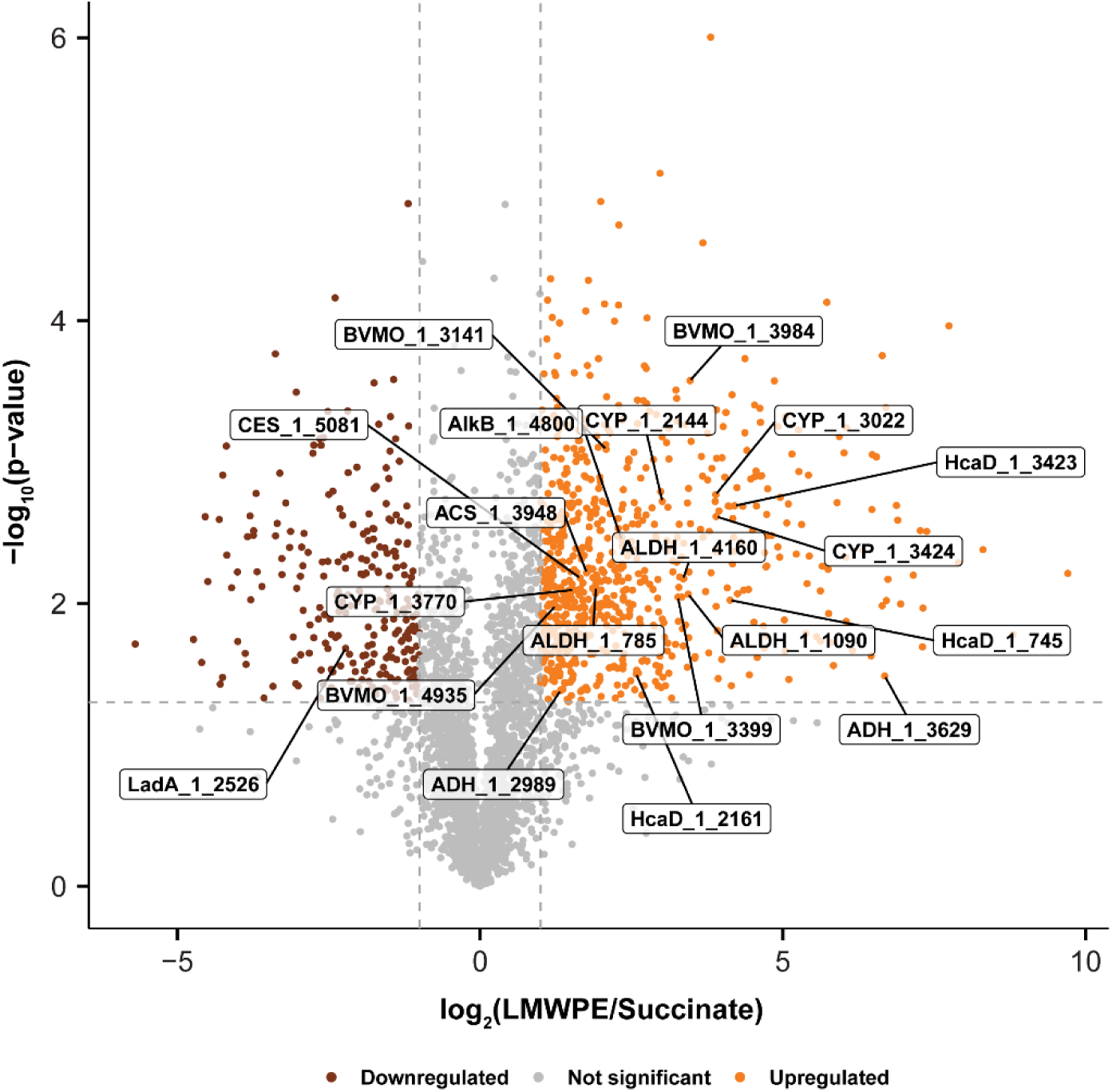
Differentially abundant proteins associated with alkane and 2-ketone degradation detected in the LMWPE-derived proteome of *Rhodococcus* sp. ASF-10. KEGG Orthology IDs, as well as BLAST hits were used to identify and highlight proteins known or hypothesized to be involved in alkane and 2-ketone degradation. The dashed horizontal line denotes the *P* value cut-off (*P* < 0.05), and the dashed vertical lines denote the log_2_-fold change cut-off (< −1 and > 1). Source data is provided in **Table S3A**.

Four predicted flavin-binding monooxygenases (FMOs) were also identified as differentially abundant in the LMWPE-derived proteome of *Rhodococcus* sp. ASF-10. FMOs belong to a family of xenobiotic-metabolizing enzymes, which can convert a diverse range of substrates. The family includes Baeyer–Villiger monooxygenases (BVMOs), enzymes involved in the subterminal oxidation of alkanes (69), where they oxidize carbonyl groups into an ester (70). We predicted the 3D structures of the putative *Rhodococcus* sp. ASF-10’s FMOs using AlphaFold3, which matched known BVMOs in the AFDB–SwissProt database. (**Figure S1**, **Table S3B**). Overall, the presence of multiple differentially abundant BVMOs (BVMO_1_3141, BVMO_1_3399, BVMO_1_3984 and BVMO_1_4935) in the LMWPE proteome of *Rhodococcus* sp. ASF-10 supports their involvement in oxidizing 2-ketones present in the substrate into esters. Among the differentially abundant proteins, a carboxylesterase (CES_1_5081) was also detected (**Figure 6**). This enzyme might be involved in the hydrolysis of esters with formation of a primary alcohol and an acetate. The primary alcohol can be further metabolized by an alcohol dehydrogenase and an aldehyde dehydrogenase into a fatty acid, while the acetate can directly enter the tricarboxylic acid (TCA) cycle (58). An acetyl-CoA synthetase (ACS_1_3948) was also more abundant in the LMWPE proteome, suggesting that acetate produced from ester hydrolysis could be converted to acetyl-CoA. (**Figure 6**). Following catabolism of the primary alcohol, intermediates can be funneled into the pathway for β-oxidation of fatty acids. Indeed, enzymes involved in the β-oxidation pathway were also differentially abundant in the LMWPE-derived proteome, namely the acyl-CoA dehydrogenase ACD_1_4907, the long-chain acyl-CoA dehydrogenase ACADL_1_1712, the enoyl-CoA hydratase EchA_1_2812, the 3-hydroxyacyl-CoA dehydrogenase FadJ_1_4040 and the β-ketothiolase FadA_1_1777 (**Table S3A**).

The levels of the enzymes involved in the pathway for 2-ketone metabolism were higher in the LMWPE proteome, yet the mechanism by which the bacterium transports the 2-ketones into the cell remains unclear. Fatty acid transport systems, such as lipid-transfer protein, or more unspecific hydrophobic transport complexes (71) or ABC transporters (72) may facilitate uptake.

While specific transporters of methyl ketones have not been identified or characterized in literature, multiple ABC transporters were significantly differentially abundant in the LMWPE proteome (**Table S3A**).

A protein (locus tag: RhASF-10_2_184) exhibiting high similarity to a LMCO (LMCO2), a laccase previously reported to be PE-oxidizing (53), was detected in the LMWPE-derived proteome of *Rhodococcus* sp. ASF-10, although it was not differentially abundant (**Table S3A**). If this enzyme was truly active on PE polymers, a shift in molar mass distribution of the higher M_w_ region of the LMWPE would be observed. Such a change, however, was not detected in our analysis (**Figure 5**).

Overall, the proteomics results align with the analytical data of the spent LMWPE particles, demonstrating that *Rhodococcus* sp. ASF-10 can metabolize alkanes and their oxidized derivatives originating from PE.

### Analysis of co-expressed proteins reveals the involvement of biofilm and biosurfactants in LMWPE degradation

To elucidate correlations between the protein abundance levels, a clustering tree was constructed using *Rhodococcus* sp. ASF-10’s LMWPE and sodium succinate-derived proteomes. Proteins showing the same expression patterns and high correlation were grouped into the same branch. Then, modules were defined by merging branches with similar expression patterns (**Figure 7A**). A total of 10 modules were defined. The module showing a significant positive correlation with the LMWPE proteome (“green module” in **Figure 7B**, Corr = 0.99, *P* < 0.01) was also the biggest module including a total of 1,556 proteins (**Figure 7C**). The proteins in this module presented positive eigengene values for *Rhodococcus* sp. ASF-10 grown in LMWPE and negative for samples cultivated in sodium succinate, indicating higher abundance in the LMWPE proteome (**Figure 7C**).

**Figure 7.**
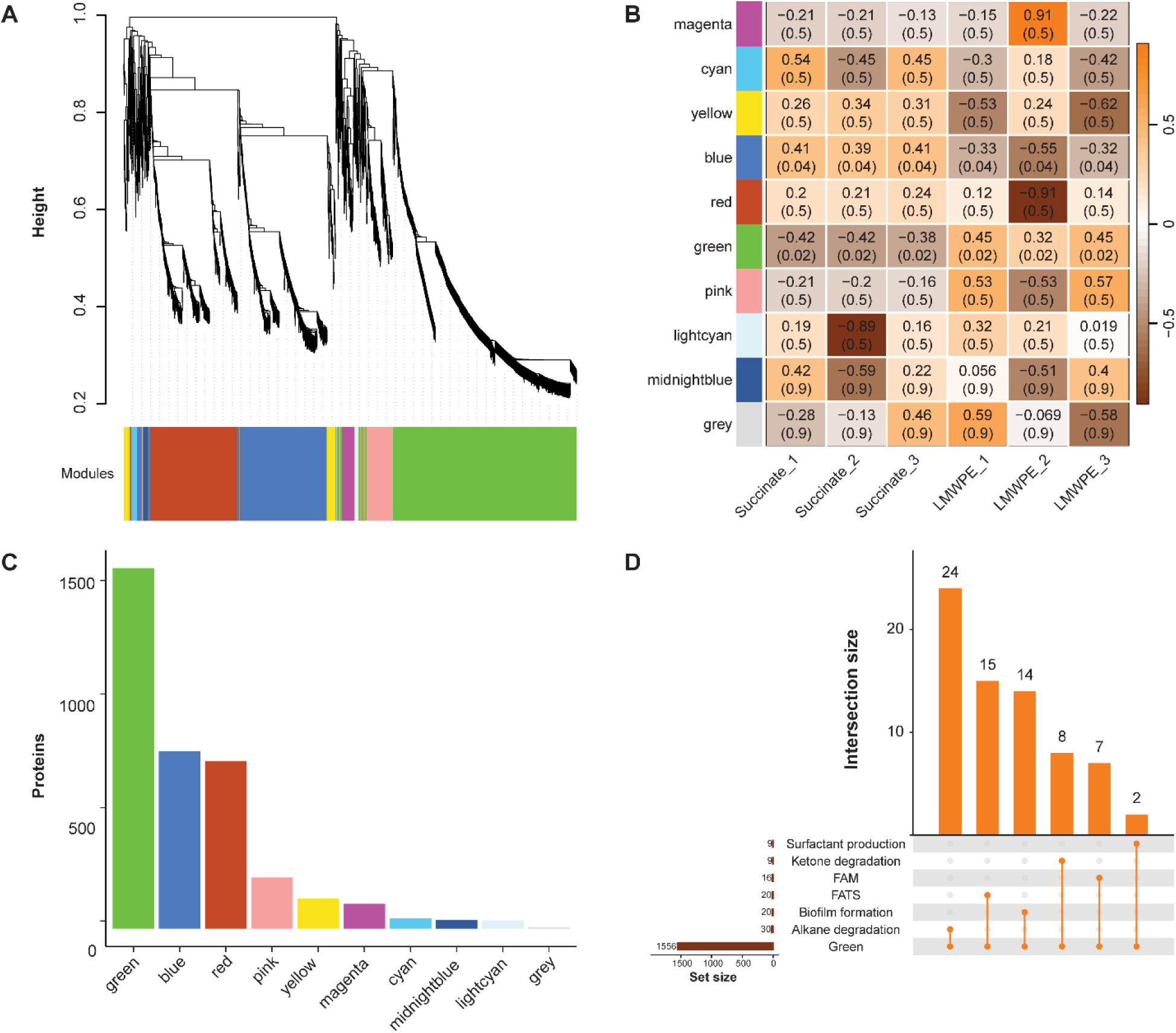
Weighted gene co-expression network analysis of *Rhodococcus* sp. ASF-10 grown in LMWPE and sodium succinate. **(A)** Cluster dendrogram of all proteins. **(B)** Distribution of protein counts per module. **(C)** Heatmap of module-eigengenes per sample. The number on top depicts the eigengene value, while the numbers between parentheses indicate the module-trait significance (*P* value). **(D)** Upset plot with the intersections between the proteins in the green module and proteins related to biofilm formation, fatty acid transporter systems (FATS), fatty acid metabolism (FAM), surfactant production, and degradation of alkanes and 2-ketones. Source data is provided in **Table S4**.

Pathways containing proteins of interest were intersected with the proteins belonging to the green module (**Figure 7D**). Specifically, this module contained proteins involved in the degradation of alkanes and 2-ketones, fatty acid transport systems and β-oxidation of fatty acids. In addition, two enzymes associated with production of surfactant and fourteen enzymes involved in biofilm formation were detected, suggesting their possible contribution in facilitating surface attachment, emulsification, and stable colonization of LMWPE particles by *Rhodococcus* sp. ASF-10. Trehalose containing glycolipids are biosurfactants produced by *Rhodococcus* spp. and other members of the Mycobacteriales Order. The production of a surfactant has been previously reported during the growth of *Rhodococcus* spp. on *n-*alkanes (73, 74). Biosurfactants can aid in substrate accessibility, particularly during the degradation of hydrophobic water-insoluble substrates (75), such as in the case hydrophobic alkanes and 2-ketones. Attachment of bacterial cells onto the LMWPE surface through biofilm formation could also aid in substrate accessibility through closer proximity to the hydrophobic PE derivatives.

Of note, the *Rhodococcus* sp. ASF-10’s LMCO (LMCO2_2_184), which is highly similar to an enzyme previously reported to mediate PE degradation (53) was not detected in the LMWPE associated module. These findings further suggest that this enzyme is unlikely to be involved in the degradation of polymeric PE, nor the PE-derivatives present in the oxidized LMWPE substrate.

### Concluding remarks

In this study, we systematically investigated the genomic basis for hydrocarbon decomposition by *Rhodococcus sp.* ASF-10 within the context of current knowledge on *Rhodococcus* spp., followed by proteomics-based identification of key enzymes employed by ASF-10 to mineralize molecules derived from LMWPE. The distillation of functional annotations into metabolic traits revealed a gradient of metabolic capacities among *Rhodococcus* spp, which correlated with their genome sizes up to 6 Mb, and were independent of the isolation source (**Figure 3**). Alkane utilization emerged as a core function among all the *Rhodococcus* genomes analyzed (**Figure 4**). In-depth analytical and proteomic insight demonstrated that *Rhodococcus sp.* ASF-10 can selectively metabolize alkanes with carbon length up to C29 and 2-ketones up to C28 when growing in LMWPE, while the polymeric fraction of this substrate remained undegraded (**Figure 5**). Proteomic analysis and WGCNA showed that this bacterium utilizes a repertoire of alkane and 2-ketone degrading enzymes, such as AlkB and BMVOs, and produces biosurfactants and biofilms in its metabolism of LMWPE (**Figure 6** and **7**). Remarkably, proteins previously indicated to be involved in PE utilization, including laccases and LMCOs, by other *Rhodococcus* isolates were not differentially abundant in our proteomic analysis. This suggests that the actual breakdown of the polymeric component by these enzymes may need to be reassessed. The importance of combining advanced substrate characterization (for instance using analytical techniques such as SEC and GC-MS) before and after bacterial growth to accurately determine whether microbes can cleave the PE backbone and subsequently mineralize it cannot be exaggerated, especially as the field of plastic degradation is increasingly searching for microbes and their enzymes to tackle this global pollution problem.

Overall, our findings provide significant insights into the capability of the salmon gut isolate *Rhodococcus sp.* ASF-10 to metabolize alkanes and 2-ketones from abiotically oxidized LMWPE and open venues to potentially exploit this microbe and its enzymes as a bioremediation tool for hydrocarbons and microplastic derivatives in aquaculture settings.

## Acknowledgements

The authors thank Eirik Degré Lorentsen and Ingrid Bakke at the Norwegian University of Science and Technology (NTNU) for providing the *Rhodococcus* sp. ASF-10 used in this study. We express our gratitude to MSc student Thea Samskott for conducting preliminary growth experiments of the isolate on various substrates. In addition, we thank Asbjørn Iveland, Bavan Mylvaganam, Sara Rund Herum, and Steffen Annfinsen at Norner AS for their valuable analytical support.

## Data availability

The genome sequence, gene annotations and predicted proteins have been made publicly available via FigShare (DOI: 10.6084/m9.figshare.30384346). The mass spectrometry proteomics data has been deposited to the ProteomeXchange Consortium via the PRIDE (Proteomics Identification Database) partner repository with the data set identifier PXD069510. All data is publicly available as of the date of publication and complies with the data reuse guidelines presented in Hug et al. 2025 (76).

## Supplementary figure

**Figure S1.**
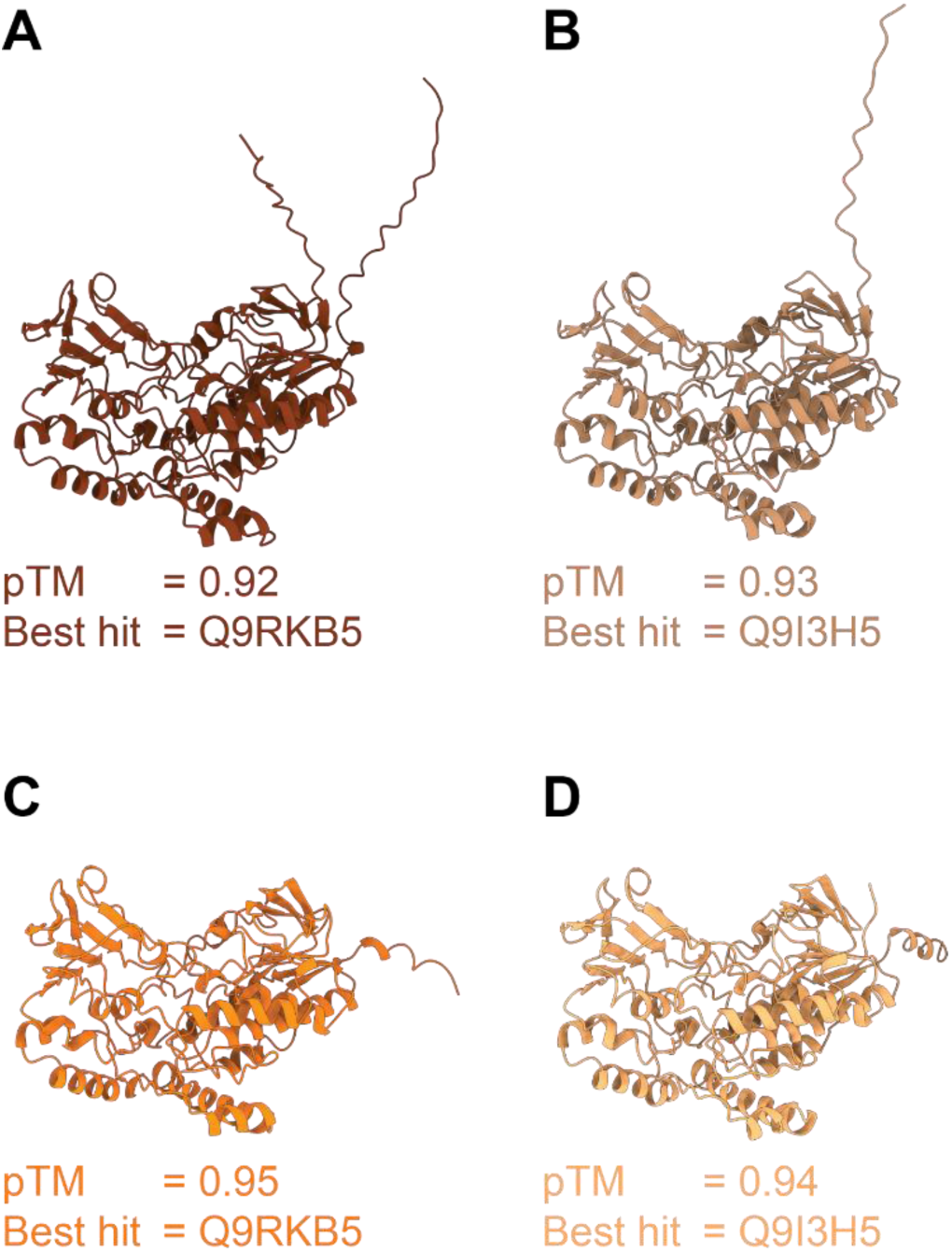
Predicted AlphaFold3 structures for the differentially abundant BVMOs detected in the *Rhodococcus* sp. ASF-10 proteomics data: BVMO_1_3141 **(A)**, BMVO_1_3399 **(B)**, BVMO_1_3984 **(C)** and BVMO_1_4935 **(B)**. AlphaFold3 structure prediction scores (pTM) and the UniProt accession number for the Foldseek best hit in the AFDB-SWISSPROT database are included in the figure, and e-value and amino acid sequence identity are provided in **Table S3B**.

